# PrimedSherlock: A tool for rapid design of highly specific CRISPR-Cas12 crRNAs

**DOI:** 10.1101/2022.09.19.508610

**Authors:** James G. Mann, R. Jason Pitts

## Abstract

**Background:** CRISPR-Cas based diagnostic assays provide a portable solution which bridges the benefits of qRT-PCR and serological assays in terms of portability, specificity and ease of use. CRISPR-Cas assays are rapidly fieldable, specific and have been rigorously validated against a number of targets, including HIV and vector-borne pathogens. Recently, CRISPR-Cas12 and CRISPR-Cas13 diagnostic assays have been granted FDA approval for the detection of SARS-CoV-2. A critical step in utilizing this technology requires the design of highly-specific and efficient CRISPR RNAs (crRNAs) and isothermal primers. This process involves intensive manual curation and stringent parameters for design in order to minimize off-target detection while also preserving detection across divergent strains. As such, a single, streamlined bioinformatics platform for rapidly designing crRNAs for use with the CRISPR-Cas12 platform is needed. Here we offer PrimedSherlock, an automated, computer guided process for selecting highly-specific crRNAs and primers for targets of interest.

**Results:** Utilizing PrimedSherlock and publicly available databases, crRNAs were designed against a selection of Flavivirus genomes, including West Nile, Zika and all four serotypes of Dengue. Using outputs from PrimedSherlock in concert with both wildtype A.s Cas12a and Alt-R Cas12a Ultra nucleases, we demonstrated sensitive detection of nucleic acids of each respective arbovirus in in-vitro fluorescence assays. Moreover, primer and crRNA combinations facilitated the detection of their intended targets with minimal off-target background noise.

**Conclusions:** PrimedSherlock is a novel crRNA design tool, specific for CRISPR-Cas12 diagnostic platforms. It allows for the rapid identification of highly conserved crRNA targets from user-provided primer pairs or PrimedRPA output files. Initial testing of crRNAs against arboviruses of medical importance demonstrated a robust ability to distinguish multiple strains by exploiting polymorphisms within otherwise highly conserved genomic regions. As a freely-accessible software package, PrimedSherlock could significantly increase the efficiency of CRISPR-Cas12 diagnostics. Conceptually, the portability of detection kits could also be enhanced when coupled with isothermal amplification technologies.

## Background

Sherlock-Hudson and DETECTR are two rapid nucleic acid detection tools which have recently been demonstrated to have high specificity and portability [1, 2, 3]. Both platforms rely on leveraging the distinct enzymatic properties of CRISPR enzymes [3]. Guided by a crRNA, complexed Cas12 or Cas13 enzymes target crRNA complimentary nucleic acids. Upon target recognition Cas12 and Cas13 enzymes undergo conformational changes activating a collateral cleavage effect [4, 5]. Coupled with fielded extraction, isothermal reverse transcription, and amplification potential nucleic acid targets can be detected via the utilization of the collateral cleavage effect with CRISPR-Cas specific reporter ssDNA or ssRNA fam-biotin reporter oligos on lateral flow test strips (Figure 1) [1, 6].

**Figure 1:**
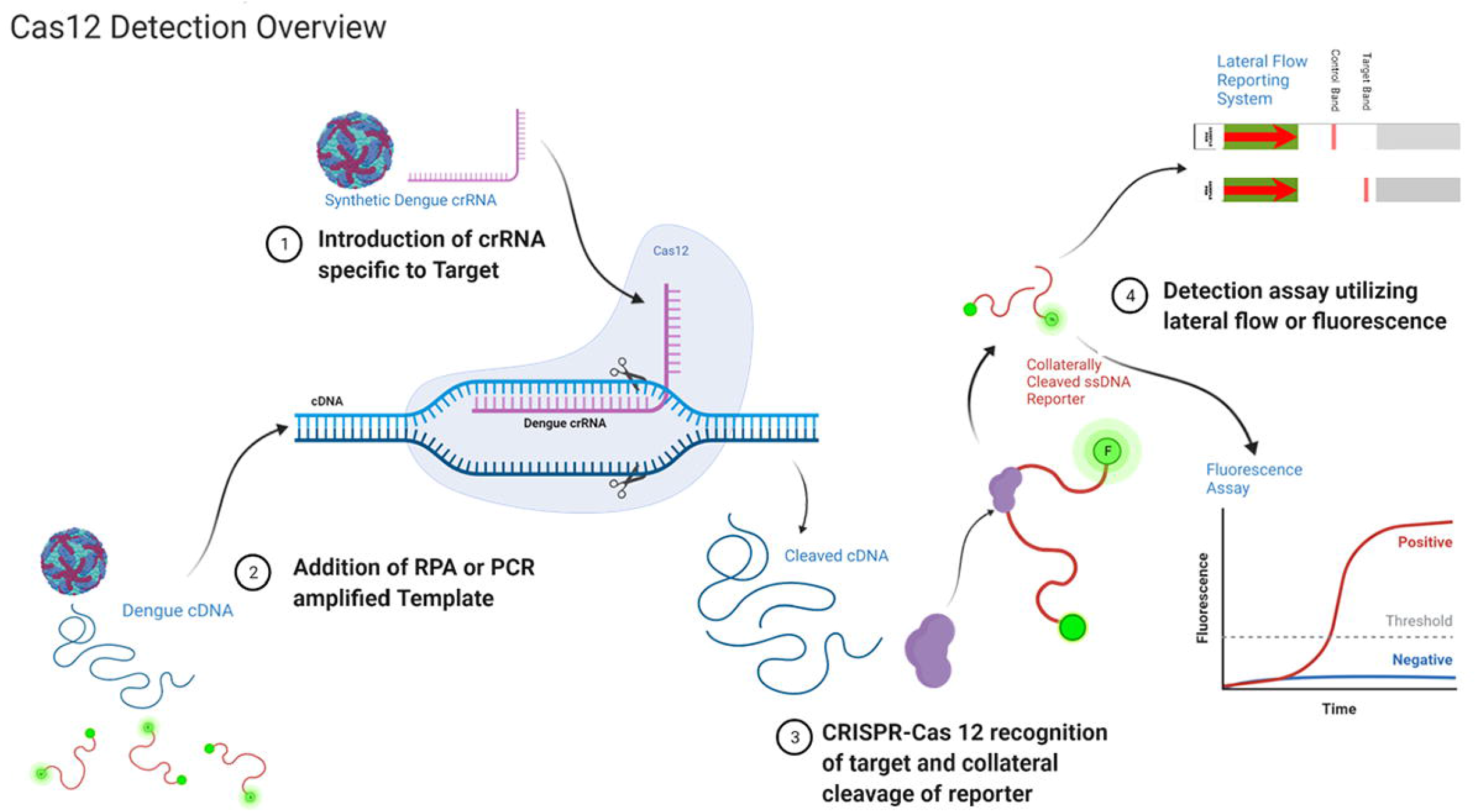
Overview of CRISPR-Cas 12 based assay approaches. Firstly, a crRNA is complexed with a Cas12 enzyme in reaction buffer. Secondly, reporter oligos and sample RT-RPA or RT-PCR product are introduced to the reaction buffer. Thirdly, target sequence recognition causes enzyme activation and collateral cleavage of reporter ssDNA. Lastly, reporter molecules either FAM-Biotin or FAM-Quencher are respectively reported on via lateral flow or Fluorescence Assay.

These assays are versatile and suitable for deployment in regions not currently accessible by industry standard techniques such as qRT-PCR. These assays have been granted FDA approval for detection of SARS-CoV-2 further demonstrating their potential as a fieldable detection assay for flaviviruses [6, 7, 8].

To date, a majority of downloadable or web-based CRISPR tools have been applied to genome editing [9, 10]. The primary function of these tools is to identify potential sites to study functional consequences of genomic, transcriptomic, and epigenomic perturbations [11, 12]. In other cases, they are used to find suitable sites for genomic inserts such as full genes or to modify existing endogenous genes [13]. In most cases, these tools have specifically focused on a single subset of potential applications utilizing CRISPR systems. When utilizing them for diagnostic purposes, many are excluded because they fail to support searches for non-standard Cas9 variants or any other CRISPR-Cas enzymes. Other tools that have expanded flexibility for user-defined PAM sequences restrict users to searching model organisms or prebuilt indices. Others, which demonstrate greater flexibility, still fail in their calculated off-target sequence tolerance. For diagnostic purposes, this caveat could lead to significantly reduced accuracy in the detection of targets [14, 15]. As yet, full-service crRNA design tools are lacking for the development of CRISPR-Cas based diagnostic platforms. Here we demonstrate the utility of an automated approach, which we have named PrimedSherlock, that is intended for CRISPR-Cas diagnostic crRNA discovery and analytical evaluation.

PrimedSherlock is a tool that can either act independently or as a companion to PrimedRPA. Users provide either the output of PrimedRPA or user-generated primer pairs for crRNA target generation. The PrimedSherlock tool then screens for ideal crRNA targets within the consensus amplicon of all genomes within the provided on-target dataset. The algorithm uses a simple design logic and revolves around two principles: firstly, crRNAs are only permitted to move forward to on-target processing if they contain more than 10 mismatches to potential near-match background genomic sequences, and secondly, mismatches are heavily penalized in the on-target screening phase. The discrimination value can be set by the user. However, the default discrimination value is three mismatches, with no regard to position along the 5’ seed region or peripheral 3’ end.

We have validated the utility of this approach by demonstrating specific detection of human disease pathogens. Among these are arboviruses in the Flaviviridae family that are transmitted by mosquitoes. More than 80% of the world’s population is currently at risk for vector-borne diseases [16]. Globally, rates of mosquito-borne illnesses are increasing, which highlights the vital need for the introduction of rapid, reliable, field detection systems. In this study, we describe the results of our design process and subsequent fluorescence detection assays. More specifically, we have demonstrated the generation of highly accurate crRNA pairs from RPA primers derived from the PrimedRPA tool. We have also validated the coverage of tool-generated crRNA pairs by utilizing non-infectious genomic RNAs from Dengue virus serotypes 1 through 4 (DENV1-4), Zika Virus (ZIKV) and West Nile Virus (WNV). We expect this tool to serve as an integral part of the crRNA design process for CRISPR-Cas based diagnostic approaches.

## Results

### Hardware benchmarking

To better understand tool performance on diverse hardware systems, we performed benchmark testing utilizing two different enthusiast setups (Figure 2). The first platform tested was an AMD 3900XT with an Nvidia 2080ti and the second consisted of a 5950X and 3080ti. Benchmarking consisted of varying the total thread count utilized for the tool (Figure 2). Changes to our user configurable thread count led to significant differences in runtime, consistent across hardware setups (Figure 2).

**Figure 2:**
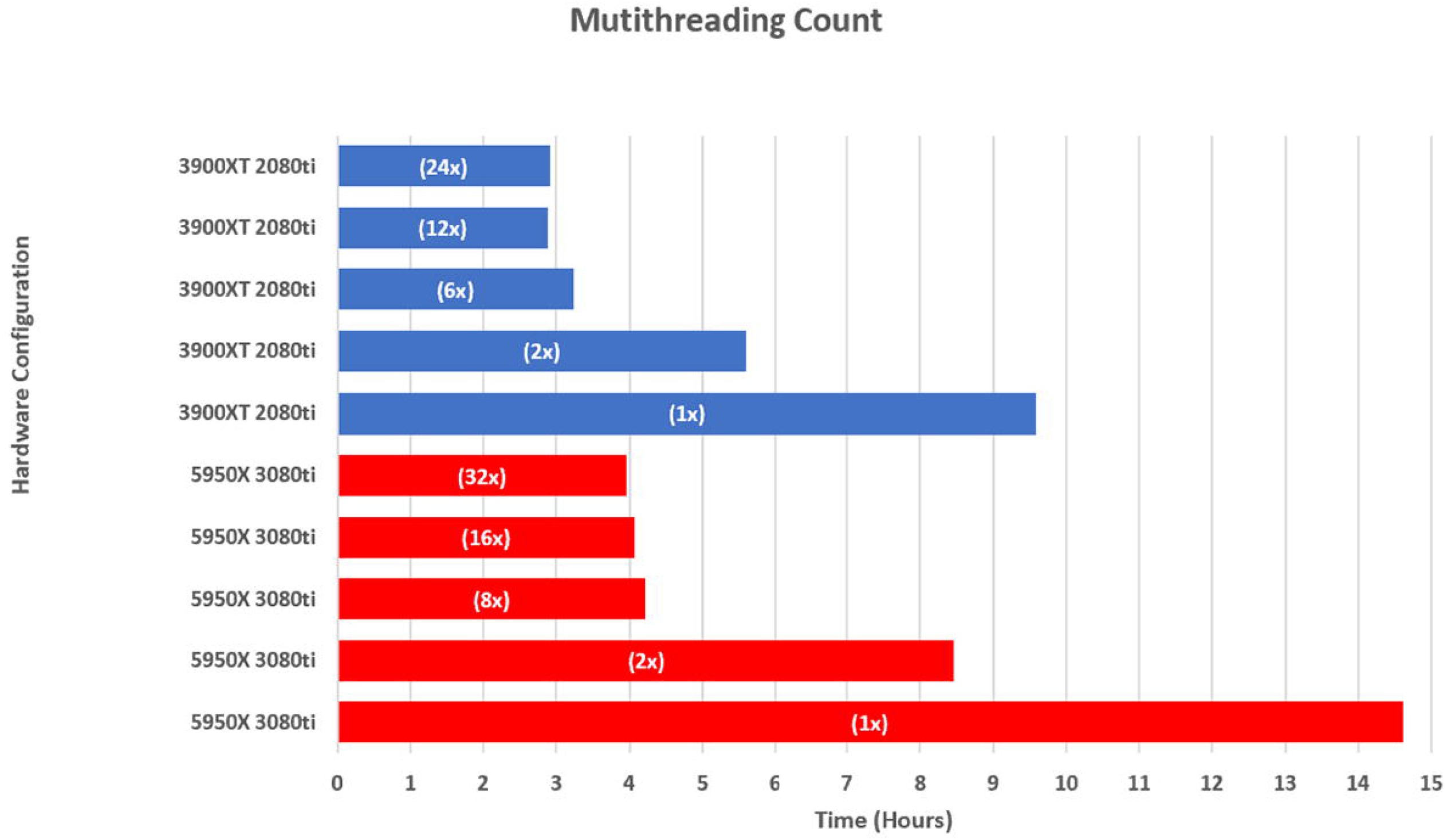
Benchmarking of varied hardware setups, configuration indicated on the left. Thread configuration indicated in bar as (thread count x). All tested configurations and the average runtime of the tool in hours dependent on core count with *thread count indicated in white*.

Running the tool with a single thread, spawning a single instance of Cas-OFFinder, led to runtimes of greater than 8 hours (Figure 2). The 5950X setup required a total of 14.6 hours while the 3900XT required 9.6 hours (Figure 2). Run configurations with the maximum available threads resulted in runtimes of 2.9 hours and 4.0 hours for the 3900XT (24 threads) and the 5950X (32 threads), respectively (Figure 2, Supplemental Figure 1).

Next, were performed a series of tests on thread count values of one-half, one-fourth, and one-sixth of total available threads for each configuration. The 3900XT achieved the fastest runtime of 2.9 hours with 12 threads, while the 5950X achieved a runtime of 4.1 hours with 16 threads, just slower than the 32-thread speed (Figure 2).

### Primer and crRNA Design

PrimedSherlock is a fast automated package which can extend existing bioinformatic platforms such as PrimedRPA to provide fully automated primer and crRNA design solely from curated genomic datasets (Figure 3).

**Figure 3:**
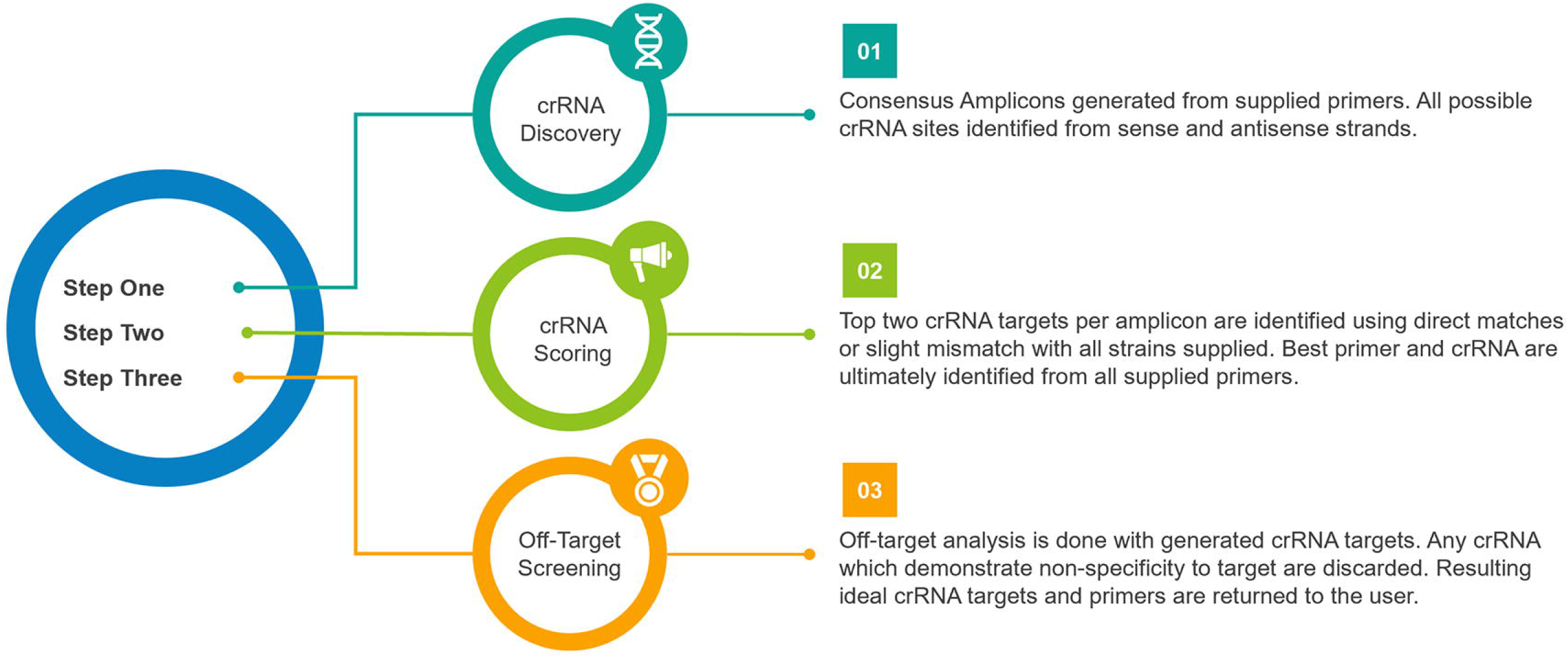
Generated primer pairs from PrimedSherlock alongside background genomic sequences and target sequences were utilized as input for PrimedSherlock. Consensus amplicons were generated to determine valid crRNA targets specific to each virus. Discovered crRNA targets were analyzed for strain coverage and potential off target recognition. Scoring was performed to determine the best two primer & crRNA pairs.

We selected six Flaviviruses for initial design and validation (Table 1). PrimedSherlock generated two sets of highly conserved primer and crRNA combinations, labeled Final_Output.csv, respectively. (Table 1 and Supplemental Files). For each pathogen, the optimum primer set and corresponding crRNA pair was selected and commercially synthesized (Table 1). As described below, crRNA targets varied in sequence conservation across viruses, and multiple crRNA targets were required for ideal coverage.

**Table 1:**
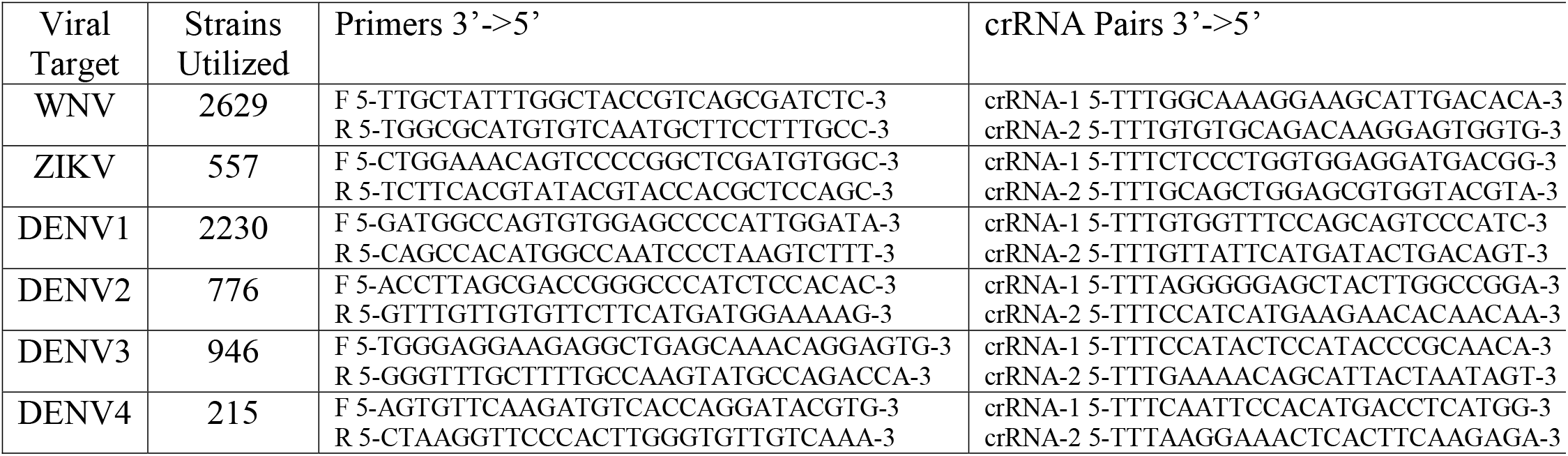
A list of the top-scoring primer pairs and associated crRNA combinations generated from PrimedSherlock. Primer pairs and crRNAs were commercially synthesized (Integrated DNA Technologies) with /Alt1/ tags added to the crRNA for stability.

For West Nile Virus, a total of 2629 genome sequences from all available strains was utilized as the input for Primer and crRNA discovery alongside a total of 6957 off-target viral strains (Supplemental Files). For our top-ranking crRNA and primer combination, 2614 strains were determined to be within our constraints for detectability. Fifteen strains were determined to have poor detectability, either possessing significant mismatches to both crRNA sequences and or possible mutation within the PAM sites. This resulted in a predicted blended crRNA target coverage of 99% of known whole genome sequences listed on NCBI GenBank (2614/2629). Our second designed set of crRNA and primer combinations had a total predicted coverage of 2523 strains (within four or fewer base-pair mismatches). However, for this set we determined that a total of 89 strains were poorly covered due to potential PAM site mutation or significant mismatches to target sequences. Our second set had a predicted blended crRNA target coverage of 95% (2523/2629).

For Zika Virus a total of 557 genomic sequences were utilized alongside 9164 off-target viral strains to develop suitable crRNA targets and primer pairs. We determined that the top-ranking crRNA and primer combination covered a total of 554 strains (with four or less base-pair mismatches). One strain, KY962729.1 was predicted to be undetectable either due to possessing significant crRNA target mismatches or mutation within the PAM. This crRNA and primer set had a predicted total coverage of 99.4% percent of full genome sequenced strains. The second generated set had a predicted coverage of 99.2% of supplied whole genome strains (553/557).

The DENV-1 primer and crRNA target design process utilized a total of 2230 full-length genomes as well as 2978 off-target serotypes or strains from the other viruses in this study. Using our analysis pipeline, we determined that 96% (2141/2230) of provided on-target strains would be detectable with the top ranked crRNA guide/primer combination, possessing four or fewer base-pair mismatches. A total of 83 strains were likely to be undetectable due to PAM mutations, and a total of six strains contained crRNA target mutations of more than four basepairs. The second ranked crRNA and primer set had a total predicted coverage of 89.1% (1987/2230) with 1987 strains possessing four or fewer base-pair mismatches for either crRNA target. A total of 224 strains had PAM site mutations or significant mismatches throughout either crRNA target.

The DENV-2 design process utilized 776 whole genome sequences alongside a total of 6206 off-target sequences. For the top-ranked crRNA and primer combination, coverage of 97.6% (758/776) of all strains was predicted, due to combined crRNA target sites possessing four or fewer base-pair mismatches for either guide. A total of 16 strains possessed PAM site mutations, and a total of two strains contained crRNA target mutations of more than four base-pairs. The second ranked crRNA and Primer set had a total of 754 strains within four base pair mismatches of either crRNA target or a total coverage of 97.4%. A total of 19 strains were determined to have mismatches in the PAM Site or significant mismatches throughout either crRNA target significantly hampering detectability.

For DENV-3 Virus a total of 946 full genomic sequences from DENV-3 diverse strains were utilized alongside a total of 4014 off-target serotype or viral strains. The top-ranking crRNA and primer combination was predicted to have a crRNA target site coverage of 100% (946/946). For the second combination, coverage of 83.8% (793/946) was predicted. A total of 144 strains possessed PAM or significant sequence mutation to likely hamper detection.

Primer and crRNA design for DENV-4 utilized 215 on-target complete genome sequences alongside 3580 off-target serotypes or viral genomes sourced from NCBI. We determined that the top crRNA/primer combination had a predicted crRNA target coverage of 97.6% (210/215), with four or fewer base-pair mismatches existing for either crRNA target sequence. Two strains demonstrated more than four mismatches and two additional strains had significant mismatches or PAM mutations rendering either crRNA site unsuitable (JN638572.1 and MN018392.1). For DENV-4, we were unable to generate a second RPA primer and crRNA set using our design tool. This was possibly due to overlap of crRNAs with off-target sequences or an inability of the algorithm to identify a secondary primer pair with suitable crRNA guides. This could represent a limitation of the tool for some targets.

### Primer Cross Reactivity

Primer design relied heavily on existing Primed-RPA features. Generated primer sets were designed with specifications where 80-90% sequence homology had to be attained across all strains and less than 65% identity for off-target serotype or viral strains had to be achieved. Primary top-ranking Primer sets were validated across a panel of synthetic gRNA and mosquito gDNA to determine potential cross amplification of undesired flavivirus or host DNA. It was determined that for our selected full sequence genomic RNA stocks that no sequenceable cross reactivity was detected for any primer set utilizing PCR or RPA. Furthermore, host genomic DNA from *Aedes albopictus* or *Ae. aegypti* did not amplify with any utilized primer set (data not shown). DENV-3 primers demonstrated low amplification of Strain-2 (BEI NR-50532) via PCR however this did not hinder down-stream assay detectability.

### Detection Assays

For all viruses or serotypes demonstrated the best crRNA and primer pairs were synthesized and utilized for detection assays of spiked gRNA individual adult female samples (Figure 4). As described previously, multiple CRISPR-Cas12 enzymes were utilized for our detection assays. For the *Acidaminococcus sp*. based assay, all viral or serotype targets were detected with crRNA targets. No off-target detection of other flaviviruses was apparent, nor amplification from mosquito genome (no template) or control samples wherein the reverse transcriptase was withheld (Figure 5). All detection assays reflected the sample level of fluorescence for each viral strain utilized, although variation in enzymatic activity existed (measured fluorescence) between the tested flavivirus samples.

**Figure 4:**
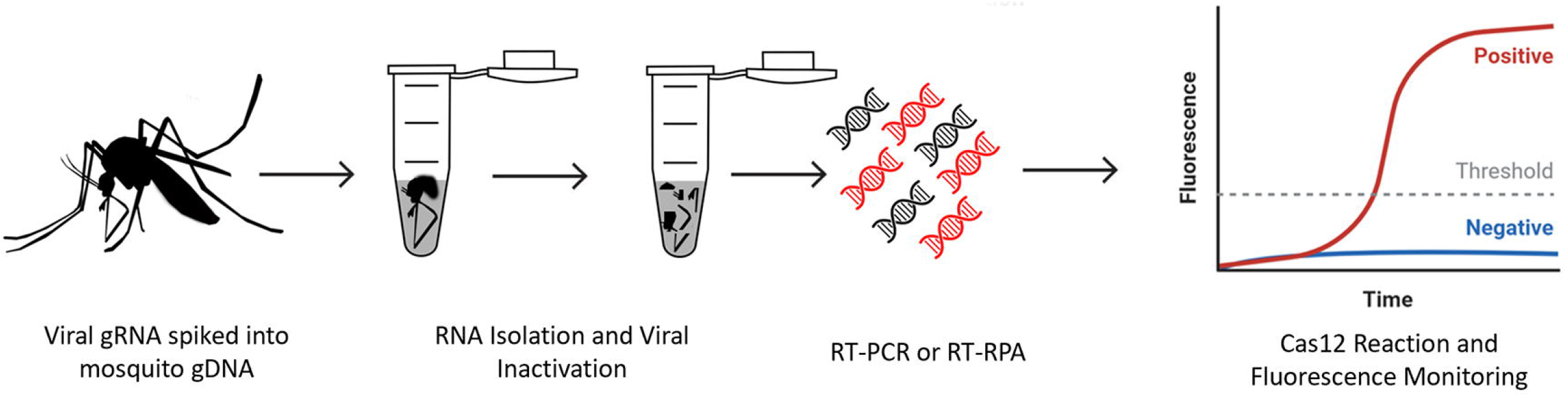
Genomic DNA from Aedes aegypti was spiked with virus genomic RNA. Samples were processed to inactivate and extract viral RNA. RT-PCR as well as RT-RPA was performed. Amplified cDNAs were utilized as templates for Cas12 fluorescence detection assays.

**Figure 5:**
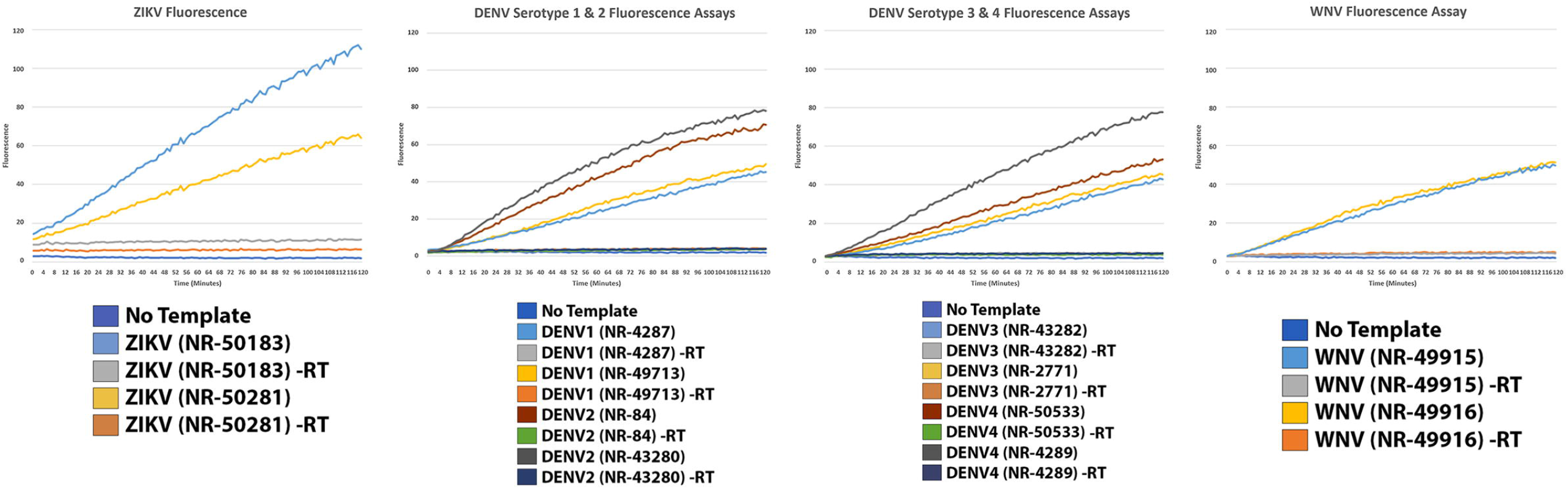
A.s CRISPR-Cas12a was used to validate PrimedSherlock crRNA targets. Fluorescence detection assays for top-ranked primer and crRNA combinations. Fluorescence values represent the average of two technical replicates measured every minute for two hours. Control assays were conducted without virus gRNA template (No Template) and for each strain without Reverse Transcriptase (-RT).

Further detection assays were performed using an altered version of *Lachnospiraceae* bacterium, called Cas12a Ultra (Integrated DNA Technologies) from separately prepared samples. All viral or serotype targets were detected without issue, with significantly reduced fluorescence activity present in all assays. Host gDNA as well as samples withheld Superscript IV reverse transcriptase demonstrated no fluorescence activity, pointing to only robust on-target activity of the enzyme (Figure 6).

**Figure 6:**
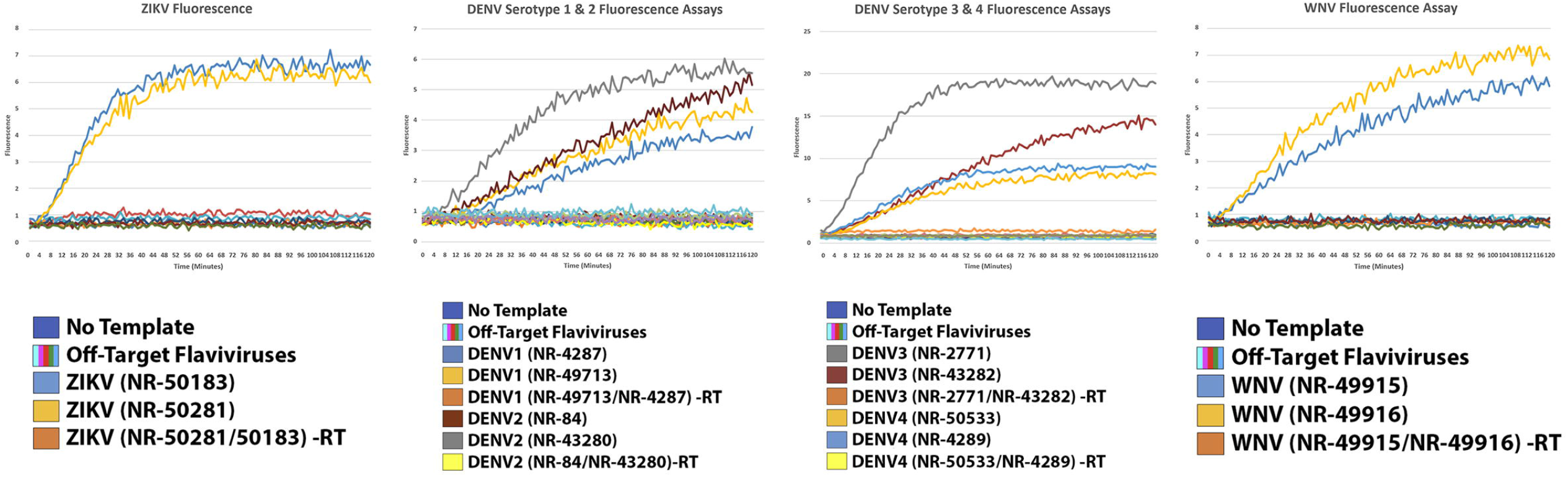
CRISPR-Cas12a Ultra was used to further validate PrimedSherlock crRNAs. Results of fluorescence detection assays for top-ranked primer and crRNA combinations. Fluorescence values represent the average of two biological replicates measured every minute for two hours. Controls assays were conducted without virus gRNA template (No Template), with two nontarget strains per sample (Off-Target Flaviviruses), and for target samples without Reverse Transcriptase (-RT).

## Discussion

CRISPR-Cas based diagnostic assays provide a platform that brings together the benefits of qRT-PCR and serological assays in terms of portability, specificity, and ease-of-use. In recent publications, both CRISPR-Cas12 and CRISPR-Cas13 have been utilized to tackle emerging viral threats [1, 2]. More recently these platforms have been rapidly shifted to aid in the detection of SARS-CoV-2, with assays utilizing both enzymes gaining FDA approval [6, 17]. The broader implementation of assays utilizing both CRISPR-Cas12 and CRISPR-Cas13 has been significantly restricted by the tedious design process required for primer and crRNA guides [18]. The design process challenges researchers to ensure their chosen targets have sequence conservation in both the chosen primer and crRNA target sequences [19]. Mismatches in the PAM sequence results in non-recognition of the target, while mismatches in the seed region decrease target recognition. Both issues lead to reduced cleavage efficiency and therefore, lowered detection sensitivity [20, 21].

In this study, we have demonstrated the utility of PrimedSherlock for designing specific crRNA guides for use with CRISPR-Cas12 platforms in conjunction with primer lists generated by PrimedRPA. For each target of interest, we leveraged publicly available genomes to design libraries of all known strain sequences as well as libraries of potential off-target viruses. In less than twenty-four hours for each target, our Python tool was able to rapidly identify and analytically evaluate the potential specificity of crRNA targets for provided primer pairs for each respective virus or serotype of interest.

The major limiting factor for the speed of the tool is the CPU and the GPU hardware present in the user’s system. Most underlying genomic analysis is powered by Cas-OFFinder. For each primer pair, any valid crRNA sequence is added to a list, which is then provided to Cas-OFFinder for on-target and off-target analysis. This is essentially a BLAST search of each on-target and off-target genome with thousands of queries of Cas-OFFinder for each crRNA. This large list of searches is divided between user-defined available threads and relies on the GPU for each individual run. Users have the option of toggling CPU-based Cas-OFFinder analysis. However, prior studies have determined that doing so leads to a 20-fold increase in run time [22], with analysis speed depending on thread count and GPU. End users can also utilize CPU-based analysis with minor code modifications that are included on the GitHub repository. In our development we utilized three diverse test systems to evaluate the rate at which ideal primer and crRNA combinations could be evaluated. The first was a 32-thread AMD 9 5950X with a founder series NVIDIA 3080TI processor. The second was a 24-thread AMD 3900XT with a founder series NVIDIA 2080TI processor. The third was a Dell XPS 15 equipped with a i7-8750H and a 1050ti processor. All tests were conducted with multithreading and utilization of all but one thread (31 threads, 23 threads, 11 threads) and GPU based Cas-OFFinder analysis. For both enthusiast level PC’s, primer and crRNA design was efficiently carried out with both PrimedRPA and PrimedSherlock with average run times of less than 8 hours for most configurations. The midrange Dell XPS took significantly longer with an average of more than two days per test run (data not shown).

The second major factor in speed is the ability for the user to diligently curate on-target and off-target datasets. For each viral or serotype target pairs of highly conserved crRNA were discovered and analytically determined to cover most strains of each pathogen listed on GenBank. However, it required manual curation of GenBank entries and elimination of rouge or mislabeled sequences. A single genome misplaced in either on-target or off-target sequences prevented PrimedRPA or PrimedSherlock from working correctly. For each program, poor curation led to false negatives with the DENV serotype datasets. The script is written in such a manner that if a crRNA target sequence is located within an Off-Target genome it is immediately blacklisted. Having a misplaced genome can result in this occurring for all target sequences, making it imperative for the user to curate the datasets in advance. For curation, we recommend using a program such as Unique Sequences, which can be found within Galaxy Tools to remove duplicate entries. We also recommend removing sequences with aberrant n counts as well as visually screening the databases for mislabeled sequences. For example, we found instances of “vaccine candidate” or “chimera” that were not viral sequences. Lastly, we recommend temporarily combining both on and off target databases and using the program Unique Seq to determine if sequences have been accidently incorporated into the incorrect database. By adopting this curation strategy, we experienced no runtime errors or issues with viral targets for both PrimedRPA and Primed Sherlock.

During refinement, we further explored the platform with several different thread count configurations of the 5950X and 3900XT setups utilizing the ZIKV dataset. We discovered that thread count has a direct effect on total runtime. Without multithreading, run times were approximately three times longer than for 24x and 32x multithreading, which were the maximum available thread counts for the 3900XT and 5950X systems, respectively (Figure 2). To our surprise, each hardware configuration reached near-minimum completion times at one-fourth the maximum thread count: 6X for the 3900XT and 8X for the 5900X (Figure 2, Supplemental Figure 1). Additionally, the 3900XT outperformed the 5950X in both single and multithreaded configurations. We believe this could be explained by the base clock speed of each CPU, which is 3.8ghz for the 3900XTand 3.4ghz for the 5900X. The 3900XT may also outperform the 5950X in per core performance. Interestingly, we found that the 12-thread configuration slightly outperformed the 24-thread configuration for the 3900XT, which could be due to minor stochastic variations in performance (Figure 2).

PrimedSherlock was originally developed as an internal tool to rapidly speed up the development of CRISPR-Cas12 assay targets. Each virus or serotype target reported within this article was designed using multiple versions of the script. Of note, the ZIKV and WNV assays were designed with earlier renditions. Earlier bench-validation efforts of analytical datasets generated by the tool provided us with valuable feedback which improved later renditions. Of important note, bugfixes to potentially issue code segments. One such bugfix was an issue with the constraints of what regions should be searched for crRNA targets. Our earliest rendition allowed for the Primer regions to be included. This resulted in one of the original crRNA targets for WNV being partially present within the primer region, which was corrected in later renditions. However, the partial presence within the primer and the formation of a primer dimer was enough to elicit a false-positive, indicating the importance of pre-screening. This phenomenon was resolved by changing the forward primer for WNV, to the one that is included in Table 1.

For any nucleic acid based diagnostic assay, a major challenge is primer conservation. Sequence mutations within the regions responsible for amplifying viral or template can easily become mutated causing reduction or complete failure in template amplification. Utilizing the Isothermal Recombinase Polymerase amplification, of particular interest to us is conservation of 5’ and 3’ ends of the forward and reverse primers. Previous studies have indicated that non-sequence homology in these regions significantly hampered the ability to amplify template [23]. In order to combat this, we set stringent conservation standards in PrimedRPA and relied on at least 80% primer sequence homology for both the forward and reverse primers across all viral targets. Utilizing two diverse strains for each viral or serotype target, we did not experience any issues with detection assays using either RT-RPA or RT-PCR based cDNA amplification nor did we experience any failed detections. As described above, we observed a significantly reduced yield of cDNA for DENV-3 Strain 2 (BEI NR-50532). We attribute this to a reduced amount of template, as compared with the other viral strains. For each NR-50532 spike-in we utilized 2ul of stock at 2.2 × 10^3^ copies per microliter. Other stocks were considerably less diluted which may resonate with the poor amplification experienced for this spike-in. However, as demonstrated in Figure 5 and Figure 6, detectability was still achieved in line with that of the other DENV-3 strain.

To validate our analytically generated crRNA pairs and primer sets we utilized two commercially available CRISPR-Cas12 enzymes. We utilized both wildtype recombinant *Acidaminococcus sp*. BV3L6 (A.s) nuclease as well as a modified recombinant *Lachnospiraceae* bacterium ND2006 (L.b) nuclease with several modifications to improve on-target editing and temperature tolerance. The choice to include both was strongly due to the influx of commercially available types, as well as manufacturer modifications and a desire to ensure usability across available CRISPR-Cas12 enzymes. For our assays we utilized multiple biological and technical replicates. For each of our included figures, each fluorescence assay graph included two averaged technical replicates representing the fluorescence detected from the detection assay. We observed consistent amplification across technical replicates. However, fluorescence varied significantly between the two versions of Cas12 tested. The apparent fluorescence reduction for the modified L.b Cas12-Ultra (Figure 6), may be due to proprietary enzyme modification(s) that may otherwise increase on-target gene editing efficiency.

Bench validation of the highest-ranking crRNA pairs utilizing fluorescence assay revealed positive detection of the target virus across all divergent strains. Although there was less fluorescence units produced by the L.b enzyme in response to target presence, there was still a considerable difference between the no template, negative RT controls, off-targets and target samples. For both, however more noticeable in the wildtype figure cleavage efficiency of the Cas12 enzyme varied by viral strain. There was a noticeable demonstration of mismatch effects being provided in Figure 5 between the two DENV-4 strains. This could either be caused by sequence variation between the crRNA site and the guide presence or the titer of each virus gDNA sample varying significantly (1.4ng/ul, NR-50533 vs 126ng/ul, NR-4289).

In drafting PrimeSherlock, we carefully considered published studies when determining the best scoring method for crRNAs. We reasoned that the most important factor for the toolset should be minimizing crRNA mismatches, especially within the PAM region, as mutations can severely disrupt dsDNA target cleavage [3]. After that, our design goal was to ensure that crRNA targets were conserved enough not to function independent of the target amplicon presence. In one study, the authors demonstrated that accumulation of mismatches lead to a diminishment of target induced off-target activity [3]. Cleavage of the non-target ssDNA was reduced and ultimately diminished at an accumulation of less than 15 base pair matches to the target nucleic acid sequence. We incorporated this constraint into the design process by excluding any crRNA possessing matches of more than 10 base-pairs to off-target genomic sequences.

In terms of crRNA specificity for target sequences, as few as two mismatches can cause a significant reduction in detection efficiency [24, 25, 26]. Furthermore, mismatches in the seed region, or bases 1-6 proximal to the PAM site, can negatively impact on-target recognition [3, 26, 27, 28]. Our design logic accounts for these mismatches to increase crRNA efficiency. PrimedSherlock is designed to select only crRNAs that demonstrate the highest conservation across all targeted strains or genomes, while mismatched crRNAs are biased against.

We further strengthened Primed Sherlock by selecting two crRNA targets within each amplicon, considering the impact of seed region and minor distal mismatches. By relying on a multiplexed approach, we greatly reduce the impact of a PAM site mutation, seed region mismatch or minor distal sequence mismatch [29]. Multiplexed approaches also reduce the need for template amplification, with larger arrays of crRNA targets diminishing the need for template amplification entirely [30]. After determining the most efficient path for crRNA design, we elected to not automate any modifications to the 3’ or 5’ ends of the identified crRNA targets, given the current debate on the effectiveness of such modifications [31, 32].

The prevention of off-target induced false positives was a major consideration during the design process. In CRISPR-Cas diagnostic assays that involve sample amplification, two factors limit the potential for off-target induced enzymatic activity. The diligent design of primers to minimize off-target amplification and the specificity of crRNAs for target sequences. For example, in assays where RPA is combined with CRISPR-Cas, either an off-target amplicon or an off-target dsDNA sequence independently wouldn’t necessarily lead to a false positive detection. However, in a multiplexed approach false positive detection in the absence of any template amplification has been documented [30]. In our opinion, combining both primer and crRNA specificity, is the best approach to reducing potential false positives. By performing off-target analysis of crRNA sequences, we can control for user-provided primer design independent of off-target enzyme activation. Further, the shift to diagnostic assays directly from sample without amplification is highly debated [33, 34, 35]. By maintaining this redundancy in our analysis, users should be able to adapt the toolset for CRISPR-Cas assays independent of sample amplification and account for the risks associated with multiplexed crRNA approaches.

An advantage of providing the original source code for PrimedSherlock is that it is modifiable as new evidence comes to light. For example, diverse nucleic acid targets may form secondary and tertiary structures, which themselves may affect the enzymatic properties of CRISPR-Cas12. Mutations in Cas12 may also alter the ability of the enzyme to interact with targeted dsDNA sequences. As more studies regarding the parameters that guide crRNA target acquisition and complexed enzyme activation are conducted, PrimedSherlock can be updated to further improve outcomes.

In validating PrimedSherlock, we focused our efforts on diverse mosquito-borne viruses of medical concern. Our examples were selected as a proof of principle demonstration that given a wide assortment of strains the tool could identify conserved regions enabling highly specific CRISPR-Cas detection assays. One strength of the toolset is that users may readily determine their own diagnostic goals. On one hand, target sequences can be narrowly selected to limit coverage to the most important circulating strains or newly disseminating strains. An example could be targeting strains of ZIKV linked to microcephaly by focusing input datasets [36]. Conversely, coverage can be maximized to all available genomes sequences for a particular pathogen of interest more generally.

## Conclusion

This tool proves versatile and allows for the rapid design of crRNA targets from curated genomic datasets. In our use case, we demonstrated the tool with several flaviviruses of major concern and further demonstrated its ability to design crRNA targets for specific serotypes of a major flavivirus. For each of our generated datasets we were able to detect only the samples which contained our viral targets. In all of our assays, we readily observed detection of strain-specific viral targets, with no detection of no-template, minus-RT, or off-target viral control samples. We expect that implementation of PrimedSherlock will not only facilitate rapid discovery of conserved crRNA targets but will improve crRNA design by identifying ideal targets that may otherwise be missed by manual curation of target sequences. As such, this process will ultimately lead to a significant reduction in the time needed to create highly specific viral CRISPR-Cas12 based diagnostics. Finally, we expect PrimedSherlock to significantly aid in the development of portable and highly specific nucleic acid-based assays for the detection of vector-borne pathogens in resource limited setting via its integration with isothermal amplification techniques and lateral flow test strips.

## Methods

### Primer design

PrimedRPA from [37] was utilized to create a list of potential Recombinase Polymerase Amplification (RPA) primers specific to ZIKV, WNV and each of the four DENV serotypes. Briefly, full sequence genomic datasets specific to each virus were sourced from NCBI-GenBank. All full-length sequences were utilized with Unique Seq on the Galaxy platform to generate a list of on-target genomic input sequences [38]. Off-target genomic datasets were generated from compiling the remaining five viruses or serotypes. On-target sequences were further manually screened and then subjected to MAFFT based alignment [39]. These MAFFT alignments were then visually inspected to rule out inclusion of chimeric or attenuated strains.

Resulting fasta files were then supplied to the PrimedRPA platform as on-target and background datasets. PrimedRPA was run with default parameters, with the option to generate probe binding sites disabled and background sequence analysis enabled. The generated Output_sets.csv, were renamed in specific format which included a viral identifier prefix on the Output_sets.csv. All input files used for PrimedRPA alongside the output_sets.csv are available as supplementary files to this publication. A list of generated primers and targets are provided in Table 1.

### PrimedSherlock

The PrimedSherlock python tool, operated by command line or in an IDE such as Spyder creates and filters multiplexed crRNA pairs for use with supplied primer combinations. An operational overview of the process is provided in Figure 3. The program utilizes a simple config file which allows the user to set various parameters including target amplicon base pair size ranges, max background identity (sourced from PrimedRPA), acceptable sequence deviation (mismatched base pairs) from background genomic sequences as well as working directory, input file and output locations. The user is then prompted to input both an “input.fna” as well as a “blast_db.fasta” which act as the on-target genomic sequences and off-target background sequences respectively. The user can provide these files with a minimum requirement of two full genome sequences per file. Providing several thousand target genomes and potential background genomes significantly increase runtime. However, this can be shortened by enabling multithreading and utilizing a mid-range graphics card such as a 2080ti.

The tool starts by first generating a consensus genome from the input.fna file. It then determines all potential crRNA sites with the needed Cas12 “TTTV” protospacer adjacent motif (PAM) contained in a consensus amplicon of provided primer combo regions. It then repeats this process for the antisense strand to achieve full coverage of the consensus amplicon for each provided primer pair. For each primer pair a separate folder is generated containing potential crRNA targets with each crRNA target written as a text file with the required input format for Cas-Offinder. The program then shifts to validating the background fasta file and removes entries with abhorrent N counts. This ensures smooth running and the removal of sequences with lesser contributions to the off-target detection. The program then utilizes multithreading to run a user defined number of Cas-Offinder instances. Cas-Offinder generates a text file containing potential matches to each crRNA pair, with a listing of how many bases in the strain/sequence match the input crRNA. This is then utilized to rank the potential off-target activity of generated crRNA. That being the cut-off point in which if a match exists, the crRNA is designated unsuitable. The program then generates a list of usable crRNA targets with low potential for off-target effects and preps them for on-target analysis. Utilizing all the input on-target genomes it determines how many genomic sequences contain exact matches to the crRNA target sequence and a few mismatches outside of the PAM region. Utilizing the same format from the off-target analysis Cas-Offinder files are made for on-target analysis. Resulting files are then used to determine how constant potential crRNA targets are across genomic sequences. The tool then determines the top two primer and two crRNA combos. For each crRNA target sequence, the program lists the genomic reference IDs with high mismatches or a PAM site mutation. It then combines this with the other crRNA to provide a consensus of potentially undetectable genomic strains. These strains are then listed in the output files to allow for a user to determine if designed crRNAs are suitable for broad usage. The same process is briefly run for the primers to determine sequence coverage and the output is provided all together in a user readable Excel file (Supplemental File 1). The design logic for the tool revolves around two main principles. Firstly, the tool only permits crRNAs to move forward to on-target processing if they contains more than 10 mismatches to potential nucleic acid sequences in background or off-target genomic regions. Secondly, it heavily discriminates against mismatches in the latter on on-target sequence analysis. This can be established as a user defined value; however, the default discrimination value is three mismatches in the target sequence, with no regard to 5’ seed region or peripheral 3’ end. The top ranking crRNA and associated primer pairs were ordered as synthetic oligos from Integrated DNA Technologies (IDTdna.com).

PrimedSherlock following the above process was utilized to generate Primer & crRNA combos for both WNV, ZIKV and each DENV Serotype. Briefly, the runtime directory was setup by extracting the PrimedSherlock Github repository. For each diagnostic assay design process, the runtime directory was cleared and prepared in a specific format for WNV, ZIKV and each DENV serotype. Any data not automatically removed was hand deleted from the runtime directory. Virus or serotype specific output_sets.csv alongside blast_db.fasta and input.fna were then uploaded to the directory. After directory setup, Spyder IDE was utilized to run the PrimedSherlock tool. A user may opt to directly use the provided batch file. The resultant Final_Ouput_Sets.csv was then utilized to pull the Best RPA primer and crRNA combo set for each virus or DENV serotype.

### Sample Preparation

Nucleic acid targets for on-target CRISPR-Cas12 detection assays were sourced from synthetic genomic RNA stocks (BEI Resources). Two microliters of viral gRNA at the concentrations supplied by BEI was used to spike 25ul *Ae. aegypti* gDNA samples. Mosquito gDNA extraction followed protocol established in [40] with a modification consisting of a substitution of rear hindleg for specimen head and thorax region. Viral gRNA spiked samples were then diluted from 25μl to 140μl for use as input template for QIAamp Viral RNA Mini Kit (Qiagen, 52904). RNA isolation and viral inactivation was performed as described in the QIAamp Viral RNA Mini Kit protocol. Isolated total RNA was then diluted into DNA / RNA Shield buffer which serves to further inactivate any potentially remaining viral particles if field collected viral samples were utilized in place of viral gRNA. Lastly, samples were heated to 90°C for 2 minutes to deactivate any remaining nucleases.

### cDNA synthesis

A total of 10μl of isolated sample RNA was used as input for reverse transcriptase-based cDNA synthesis. Utilizing either Superscript IV or Superscript III manufacturer protocol was followed with slight modification (ThermoFisher). Modifications included lengthening of the annealing step from 10 minutes to 1 hour and utilization of 1ul of Random Hexamers 200μM. No substantial difference in downstream detectability was observed for substitution to either Superscript Enzyme.

### Detection Assay

First a PCR step was performed on derived mosquito pool or synthetic viral genomic cDNA. This consisted of 5μl of cDNA template, 5.5μl of RNase / DNase free H_2_O, 1μl of each RPA Primer (10μM) and 12.5μl of Green Taq Polymerase. Performed in a Fisher MiniAmp, cycling conditions consisted of 5 minutes at 95°C, followed by 40 cycles of 94°C 1 minute, 60°C 1 minute, 72°C 1 minute, then finally 72°C for 10 minutes. Amplification of cDNA was quantified via gel electrophoresis, 2% agarose gel, with 10ul of sample per lane. Following the protocol established in [20] fluorescence-based detection was performed for a panel of crRNA targets respective to each virus. Detection assays utilized 2μl of PCR product as input for 2x scaled reactions. Control reactions consisted of pools of all off-target viruses or serotypes, as well as, negative reverse transcriptase controls, and no Cas12 enzyme (Figure 5, Figure 6). Each assay utilized the multiplexed crRNA approach consisting of the two-crRNA determined to be best via PrimedSherlock.

## Declarations

### i) Ethics approval and consent to participate

Not applicable

### ii) Consent for publication

All authors have read and approved the final manuscript.

### iii) Availability of Data and Material (ADM)

All datasets generated and/or analyzed during this study are included in this published article and its supplementary information files. The tool is publicly available on GitHub under user JamesGerardMann. Direct link https://github.com/JamesGerardMann

### vi) Authors’ contributions

JGM conceived the original study with recommendations from RJP. JGM wrote scripts and conducted experiments. RJP supervised experiments. JGM and RJP analyzed data. JGM drafted the manuscript and prepared figures. JGM and RJP revised the manuscript. All authors approved the final version of the manuscript.

## vii) Acknowledgements

The authors acknowledge Viktor Zarev, Hanna Bradford, and members of the Arthropod Sensory and Neuroethology Lab for technical assistance. The authors thank Dr. Michelle Nemec and the Baylor University Molecular Biosciences Center for the use of core facility equipment. We also acknowledge Mathew Higgins for the development of PrimedRPA and his willingness to allow us to use several code segments in this tool and the Cas-OFFinder development team for providing permission to package the executable within our GitHub repository. Figures 1–3 and Supplemental figures made with BioRender. The following reagents were obtained through BEI resources, NIAID, NIH: NR-50434, NR-50284, NR-50241, NR-50433, NR-50530, NR-4287, NR-50531, NR-4288, NR-50532, NR-2771, NR-4289, NR-50533.

