## Supplementary figures and images for "PrimedSherlock: A tool for rapid design of highly specific CRISPR-Cas12 crRNAs"

### Supplementary Figure 1

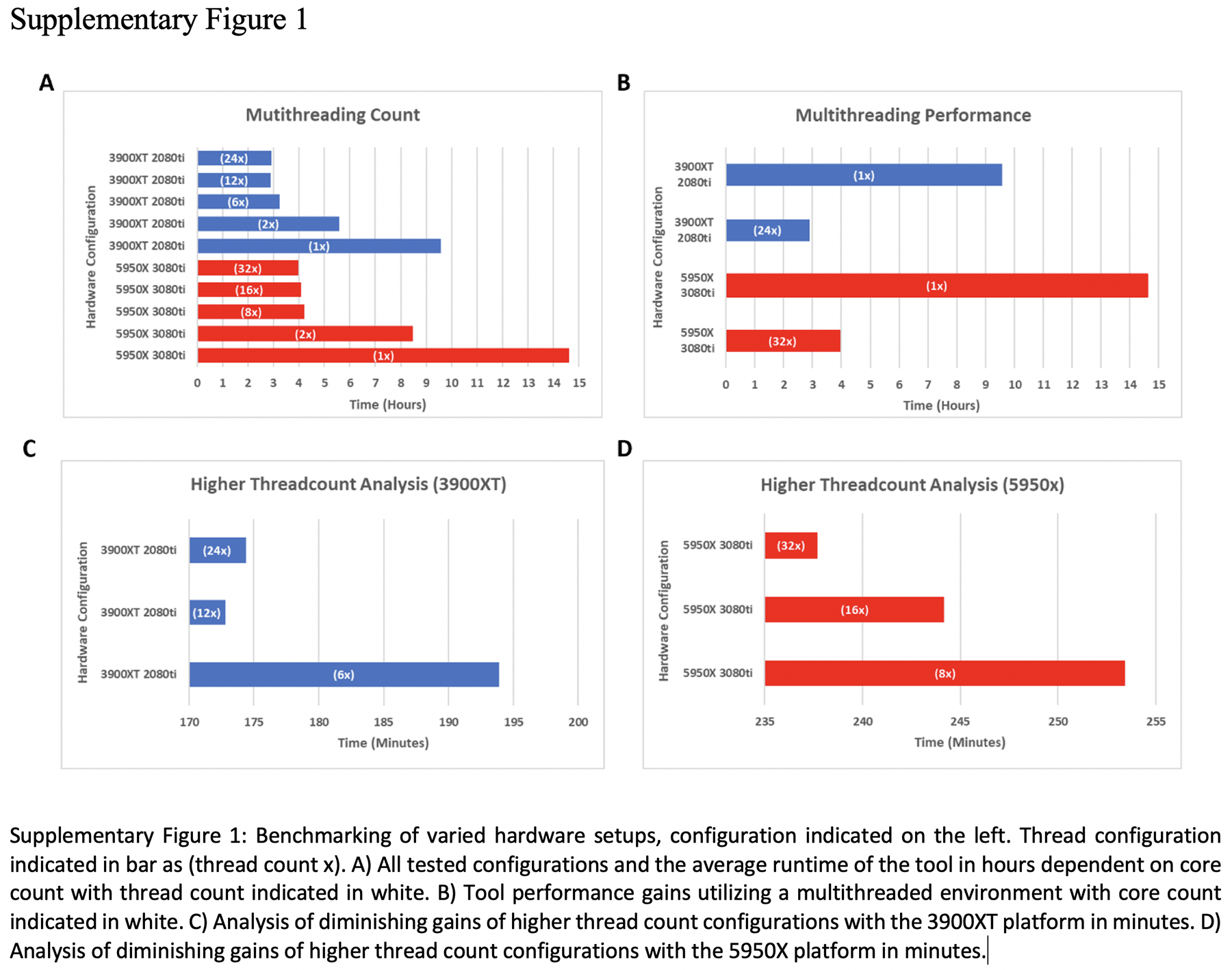
